# Insights from multidimensional analyses of post-stroke fatigue

**DOI:** 10.1101/2025.10.21.683646

**Authors:** Suhrit Duttagupta, Lutzi Castano, Sandra Chanraud, Igor Sibon, Sylvie Berthoz

**Author notes:** Corresponding author: Sylvie Berthoz, INCIA, Bâtiment Bordeaux Biologie Santé (BBS), 2, rue du Dr Hoffmann Martinot, 33076 Bordeaux CEDEX, France.

## Abstract

**Background:** Post-stroke fatigue (PSF) is an overlooked and debilitating condition. As a multidimensional construct, fatigue encompasses physical, cognitive, and emotional components, complicating efforts to understand PSF pathophysiological mechanisms and identify key predictors.

**Objectives:** We aimed to investigate the impact of lesion characteristics on the different facets of PSF while accounting for socio-demographic, psychological, and neurological factors.

**Methods:** 231 first-ever ischemic stroke patients from a prospective hospital-based cohort were assessed using the Multidimensional Fatigue Inventory (MFI) and the Hospital Anxiety and Depression Scale (HAD) alongside routine clinical evaluations. Lesion analysis was done through two approaches: a voxel-based method using support vector regression-based multivariate lesion-symptom mapping (SVR-LSM), and a network-based method using principal component analysis (PCA) of lesioned gray and white matter regions.

**Results:** The overall prevalence of PSF was 20.8%. PSF was more frequent among women and younger patients and strongly associated with HAD scores. SVR-LSM identified an association between lesions in the right corona radiata and external capsule and total MFI scores but none with HAD scores. The network-based approach showed associations between mental fatigue and reduced activity subdimensions and brain components involving cerebro-cerebellar tracts.

**Conclusions:** Our findings suggest that PSF arises from an interplay of socio-demographic, emotional, and cerebral risk factors, accounting for its heterogeneous presentation. Regarding the associations with the lesioned regions, the involvement of motor pathways raises the possibility that neuronal overactivity, compensating for disrupted networks, may contribute to long-term fatigue. Further whole-brain analyses are warranted to confirm and extend these observations.

## Introduction

Stroke is one of the main causes of reduced disability-adjusted life years (DALYs) worldwide^1^. In addition to physical disability, frequent but less apparent complications such as fatigue, emotional problems, and cognitive impairments significantly influence post-stroke outcome. Post-Stroke Fatigue (PSF) is commonly defined as “a feeling of exhaustion, weariness or lack of energy that can be overwhelming, and which can involve physical, emotional, cognitive and perceptual contributors, which is not relieved by rest and affects a person’s daily life”^2^. PSF is a frequent, long-lasting invisible condition with an estimated overall prevalence of 48%^3^. Although pre-stroke fatigue, sleep disorders, and stroke severity have been established as prominent PSF risk factors, several others remain to be considered: from sociodemographic factors, age has not been robustly associated with PSF and although female sex has been more consistently associated^4^, this may be partially explained by the tendency of women to present more severe strokes^5^. Variability in the sample characteristics such as sex ratio and stroke severity may cause discrepancy in findings across studies.

Recent meta-analyses indicated a higher prevalence of PSF following intracerebral hemorrhage^3^ while no robust association was found with neuroanatomical markers such as the extent of white matter hyperintensities or number of microbleeds^6^ - a finding recently confirmed in the CADASIL cohort^7^. Additionally, there was no robust evidence for an association between PSF and lesion characteristics such as infarct location, lateralization, or volume^3,8^. Although lesions in the brainstem and thalamus have been highlighted, it is difficult to draw firm conclusions due to substantial methodological heterogeneity^3^. Specifically, studies vary on the level of analysis: from voxel-based, regional, or network-based. Considering the inconsistent findings across these analyses, applying them based on the same cohort would provide evidence for their complementarity.

Ultimately, the potential impact of comorbid physical or mental conditions on PSF remains overlooked. In particular, although negative affectivity and fatigue share certain facets and frequently appear to be concomitant, too few studies have been conducted to strongly establish the occurrence of PSF independent of mood status^4^. Of note, while fatigue is recognized to be a multidimensional construct^9^, most studies have evaluated PSF with the unidimensional Fatigue Severity Scale (FSS). This would likely result in patients having specific domains of fatigue being undertreated.

With the current status of the literature, the present study used a multidimensional fatigue screening tool to investigate risk factors of PSF. To specifically examine stroke-related mechanisms, we recruited participants with minor ischemic strokes and no recent thymic disorders while taking potential confounding factors across physical, psychological, and clinical domains. To determine neuroanatomical risk factors, we performed lesion-symptom mapping through a voxel-based and a network-based approach.

## Materials and Methods

### Participants

The study participants were part of a larger prospective cohort recruited in Bordeaux, France (Protocol: MOTIVPOSDEP, Clinical Trials: NCT04043052) from September 2020 until September 2023 if fulfilling the following criteria: recent ischemic stroke (≤15 days ago), no severe handicap (modified Rankin scale, mRS ≤3), no post-stroke severe cognitive impairment (Montreal Cognitive Assessment, MoCA ≥16) or severe aphasia (National Institute of Health Stroke Scale, NIHSS item 9 ≥2), discharged to home, no psychoactive drug use during the preceding month, and no current or recent (over the past 6 months) mood or anxiety disorders or severe substance use disorders (except tobacco) according to the Mini-International Neuropsychiatric Interview. The study was conducted in accordance with the Declaration of Helsinki and the Institutional Review Boards and local ethics committee (CPP Sud-Est III) approved the study protocol. Written informed consent to participate and for publication was obtained from all participants prior to inclusion.

### Evaluations

Clinical assessments collected before discharge (T1) and at 3-month post-stroke (T2) included: the mRS, NIHSS (upon admission for T1), alcohol consumption and smoking dependency (Heavy Smoking Index, HSI), MoCA, and the Hospital Anxiety and Depression scale (HAD-A and HAD-D) for mood status. For both HAD subscales, a score > 7 was used to define groups with anxiety or depression. Missing data (seven HAD-A values at T1 and one at T2) were not imputed.

PSF was assessed with the Multidimensional Fatigue Inventory^9^ (MFI), a 20-item self-report questionnaire including five subscales: General fatigue, Physical fatigue, Reduced motivation, Reduced activity, Mental fatigue. Higher scores reflected greater fatigue. For the MFI total score, a cut-off of 60 was used to indicate overall PSF and the corresponding groups of participants (PSF^**+**^ versus PSF^**-**^). For the MFI subscores, the following cut-offs were used: 11 for General fatigue, 10 for Reduced activity, and 9 for Physical fatigue, Reduced motivation and Mental fatigue^10^.

### Images acquisition

Clinical MRI scans were acquired upon hospital admission with a SIEMENS Magnetron Area 1.5 Tesla scanner. Acquisition parameters for the diffusion-weighted imaging (DWI) sequence were: number of directions 3, b-value 1000, TR 5000 ms, TE 78 ms, flip angle 90°, slice thickness 2 mm, slice gap 2.4 mm, voxel size 0.78×0.78×2.4 mm^3^, field of view 270×245 mm^2^, pixel bandwidth 1455, and number of averages 8.

### Lesion preprocessing

Ischemic lesions were delineated from DWI images with the Acute-Stroke Detection Segmentation (ADS) toolbox^11^. Lesions were inspected individually and manually corrected if required using ITK-SNAP 3.8 (www.itksnap.org). Spatial normalization was performed with the BCB-toolkit (https://storage.googleapis.com/bcblabweb/index.html). Normalized lesions were resampled to a voxel size of 2×2×2 mm^3^ using trilinear interpolation. Lesion volumes were derived from normalized images to account for variability in brain volume.

### Statistical Analyses

Group comparisons were performed using the χ^2^ test of independence or Fisher’s exact test for categorical variables. Non-parametric tests were conducted as appropriate: univariate comparisons using the Wilcoxon–Mann–Whitney or Kruskal–Wallis tests with associated post-hoc analyses and effect sizes, multivariate comparisons using Quade tests, and dimensional analyses using Spearman’s simple and partial correlations. Statistical significance was set at p < 0.05. A Bonferroni correction was applied to control for false discovery rate (FDR). All analyses were performed using SPSS version 21.0 (IBM Corp.).

For neuroimaging analyses, we initially implemented multivariate support vector regression lesion-symptom mapping (SVR-LSM). Segmented lesions of the patients were inputted in the DeMarco and Turkeltaub’s toolbox^12^ (on MATLAB_R2018a, relying on the machine learning library LibSVM and on SPM12 for image manipulation), along with the patients’ MFI scores. The lesion overlap threshold was set as 5, with 1000 permutations and a clusterwise threshold of p <0.05. With corrections based on nuisance factors and clustering, the toolbox provides a voxelwise map of statistical significance. Localization of the significant clusters of voxels was done using the JHU-ICBM atlas of white matter tracts (WM)^13^.

For the MFI subscores, this method yielded no results due to the sparse overlap of lesions. Therefore, following on previous work^14^, we designed a complementary approach based on Principal Component Analysis (PCA) to reduce the dimensionality of the data while retaining as much variance as possible, taking into account the number of voxels belonging to the lesion^15^ and their location based on the Harvard-Oxford atlas for gray matter (GM) (https://identifiers.org/neurovault.collection:262) and on the JHU-ICBM atlas for WM. Using parallel analysis with Promax rotation, we identified significant loadings of regions of interest (ROIs) on distinct factors, resulting in components comprising brain regions with varying lesion volumes. These components reflected patterns of lesion distribution across patients. We then correlated the different factorial scores of the components to the scores of fatigue and clinical factors.

## Results

### Population description

Table 1 displays the descriptive statistics of the 231 participants (men/women: 163/68) at both timepoints. At inclusion, 41.1% endorsed a HAD-A score indicating probable anxiety while 7.4% endorsed a HAD-D score indicating probable depression. At 3 months, 30% endorsed a HAD-A score suggesting probable anxiety, 15.7% endorsed a HAD-D score suggesting probable depression, and 20.8% endorsed an MFI total score indicating overall PSF. The prevalence rates by MFI subdimensions were 37.7% for General fatigue, 41.6% for Physical fatigue, 31.2% for Reduced Motivation, 36.4% for Reduced Activity, and 27.7% for Mental fatigue. Over time, alcohol consumption, HSI, and NIHSS scores significantly decreased (respectively p=0.007, p=0.017 and p<0.001), while HAD scores significantly increased (both p<0.001).

**Table 1.**
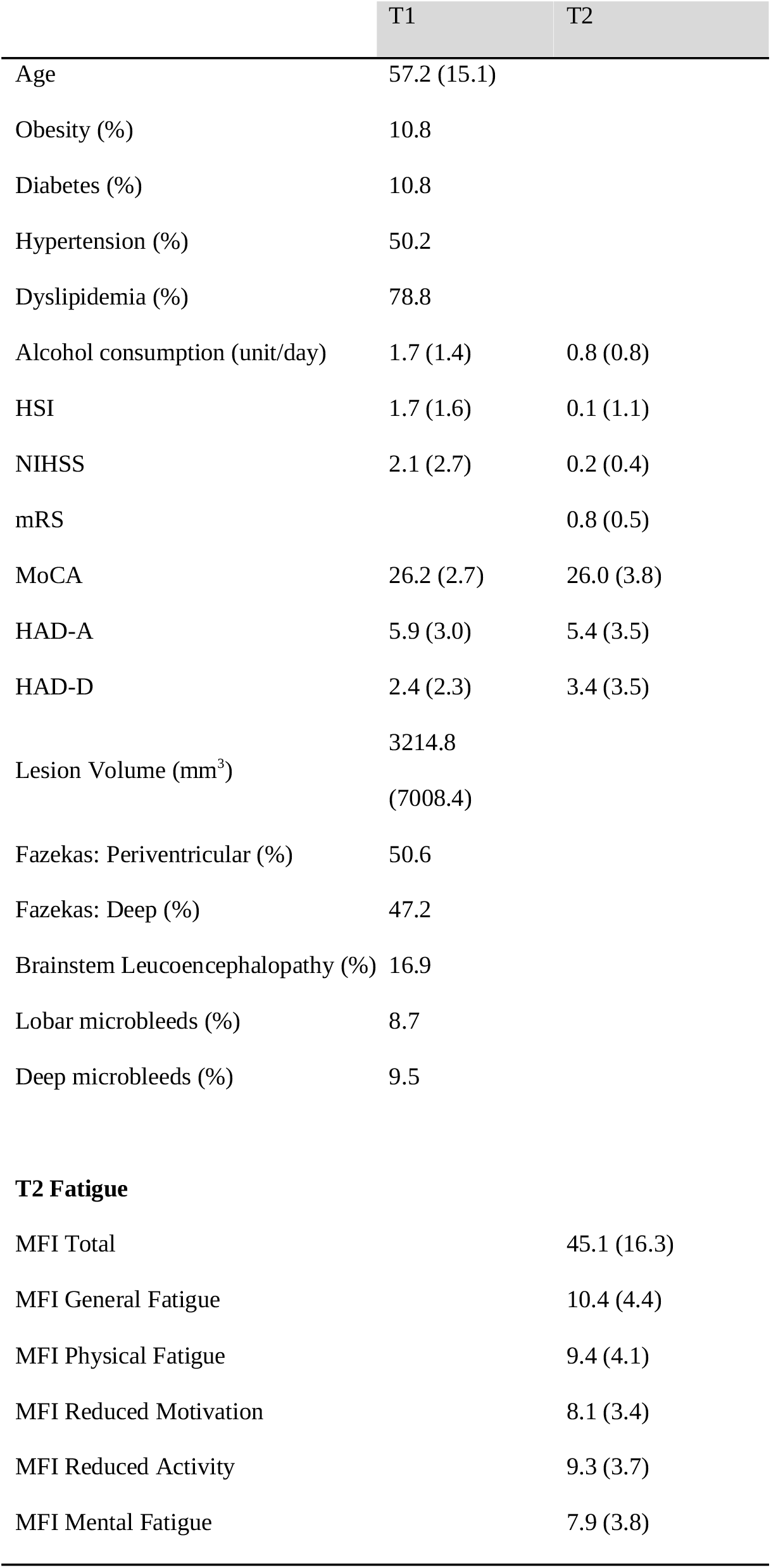
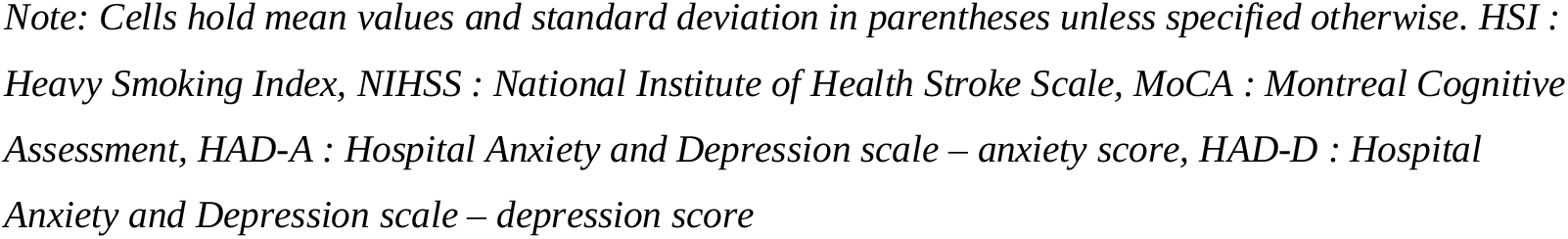
Study population characteristics at inclusion (T1) and 3 months (T2)

### Associations with socio-demographics, psychological, neurovascular and neurological factors

Univariate analyses of stroke risk factors with PSF^+^/PSF^-^ groups are presented in Table 2. The proportion of men to women was significantly different only for General fatigue (Men/Women: 32.5%/50.0%, p=0.016). Age and Mental fatigue were significantly negatively correlated (ρ= -0.205; p<0.001). The other vascular risk factors did not significantly differ by MFI groups, except for HSI, where the Motivation PSF^+^ group presented higher HSI. Univariate analyses based on sexes and the associations of potential confounding factors are presented in the supplementary material (Supplementary Tables 1-3).

**Table 2.**
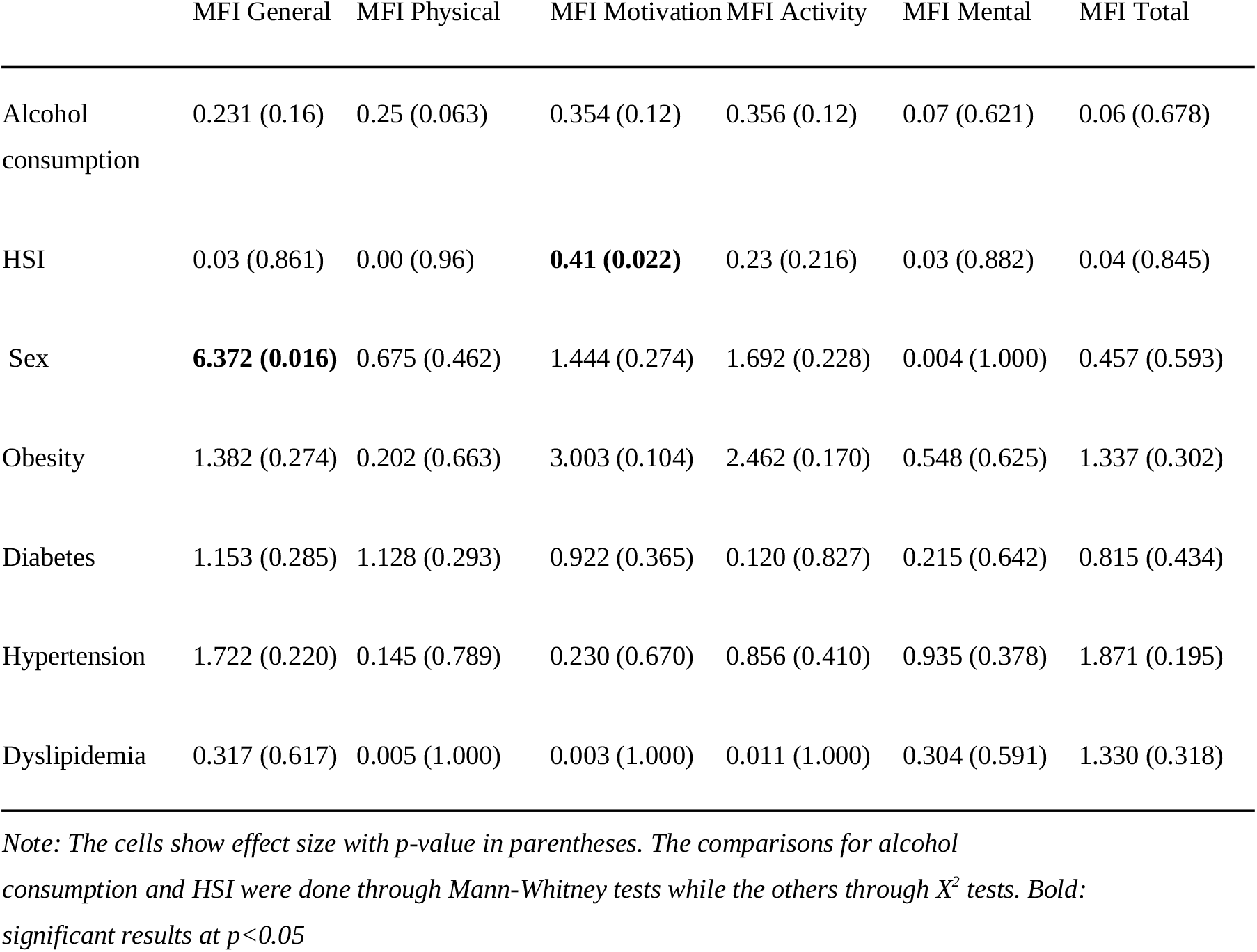
Comparison of stroke risk factors at inclusion by fatigue groups (PSF^+^/PSF-).

|  | MFI General | MFI Physical | MFI Motivation | MFI Activity | MFI Mental | MFI Total |
| --- | --- | --- | --- | --- | --- | --- |
| Alcohol consumption | 0.231 (0.16) | 0.25 (0.063) | 0.354 (0.12) | 0.356 (0.12) | 0.07 (0.621) | 0.06 (0.678) |
| HSI | 0.03 (0.861) | 0.00 (0.96) | **0.41 (0.022)** | 0.23 (0.216) | 0.03 (0.882) | 0.04 (0.845) |
| Sex | **6.372 (0.016)** | 0.675 (0.462) | 1.444 (0.274) | 1.692 (0.228) | 0.004 (1.000) | 0.457 (0.593) |
| Obesity | 1.382 (0.274) | 0.202 (0.663) | 3.003 (0.104) | 2.462 (0.170) | 0.548 (0.625) | 1.337 (0.302) |
| Diabetes | 1.153 (0.285) | 1.128 (0.293) | 0.922 (0.365) | 0.120 (0.827) | 0.215 (0.642) | 0.815 (0.434) |
| Hypertension | 1.722 (0.220) | 0.145 (0.789) | 0.230 (0.670) | 0.856 (0.410) | 0.935 (0.378) | 1.871 (0.195) |
| Dyslipidemia | 0.317 (0.617) | 0.005 (1.000) | 0.003 (1.000) | 0.011 (1.000) | 0.304 (0.591) | 1.330 (0.318) |

Note: The cells show effect size with *p*-value in parentheses. The comparisons for alcohol consumption and HSI were done through Mann-Whitney tests while the others through $X^2$ tests. Bold: significant results at $p < 0.05$

Multivariate comparisons of PSF groups are presented in Supplementary Tables 4 and 5. Results that remained using an FDR correction or adjustment for potential confounders were:

- More women than men were categorized as having General fatigue;
- Younger age was associated with having Mental fatigue;
- Higher NIHSS scores were associated with having Physical fatigue;
- Higher HAD-D scores at T1 and T2, as well as higher HAD-A scores at T2 were associated with having overall PSF and all fatigue subdimensions.

### Associations with lesion location

VLSM analysis identified a correlation between MFI Total score and lesions located in the right corona radiata and external capsule (Figure 1). The potential confounding effect of HAD-D or HAD-A scores was ruled out since no significant clusters were associated with these scores.

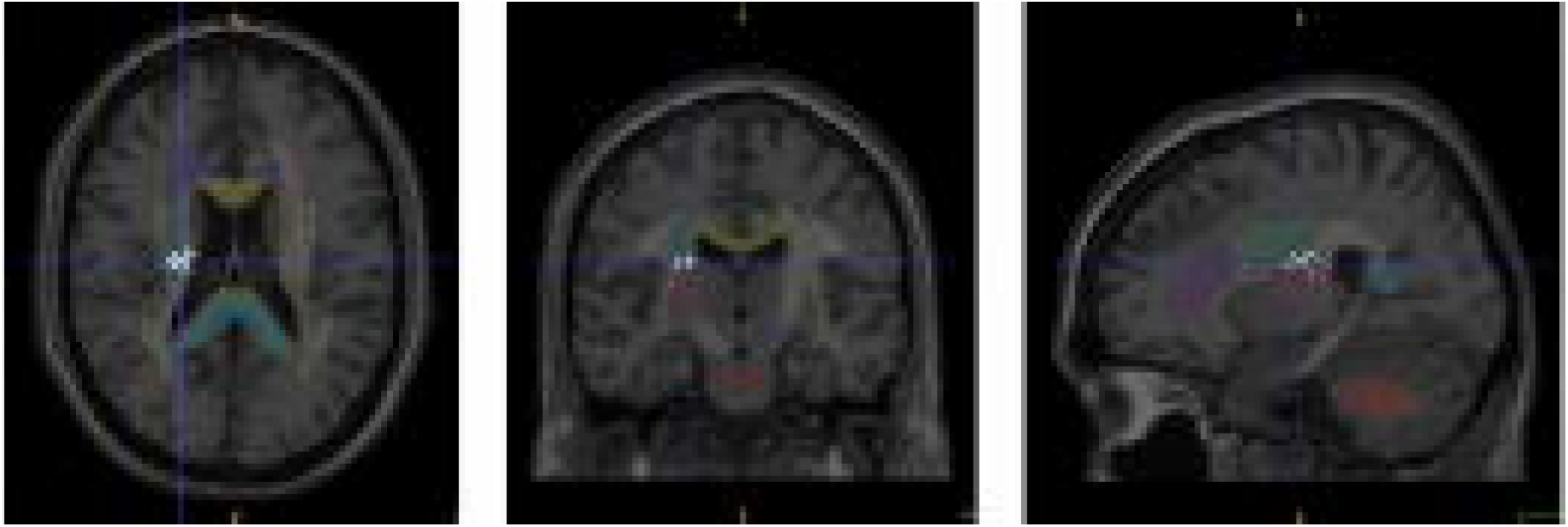

Regions associated with the MFI subdimensions could not be captured by this approach because of limited overlapping lesions. Therefore, we opted for a network-based approach based on an application of principal component analysis (PCA)^13^. Since we obtained a cluster in the white matter (WM) significantly associated with MFI total score, we based our analyses on WM regions to deepen our results as well as gray matter (GM) regions to explore what might not have been captured by the SVR-LSM analysis.

Using PCA, 12 brain components of sets of WM regions and 11 brain components of sets of GM regions were extracted (Supplementary Figure 1, Supplementary Table 6). Overall, when adjusting for potential confounders and applying FDR correction, one component (WM5) including the medial lemniscus and the cerebellar peduncles (Figure 2) was significantly correlated with Reduced Activity (ρ=0.216; p=0.001) and Mental fatigue (ρ=0.202; p=0.002).

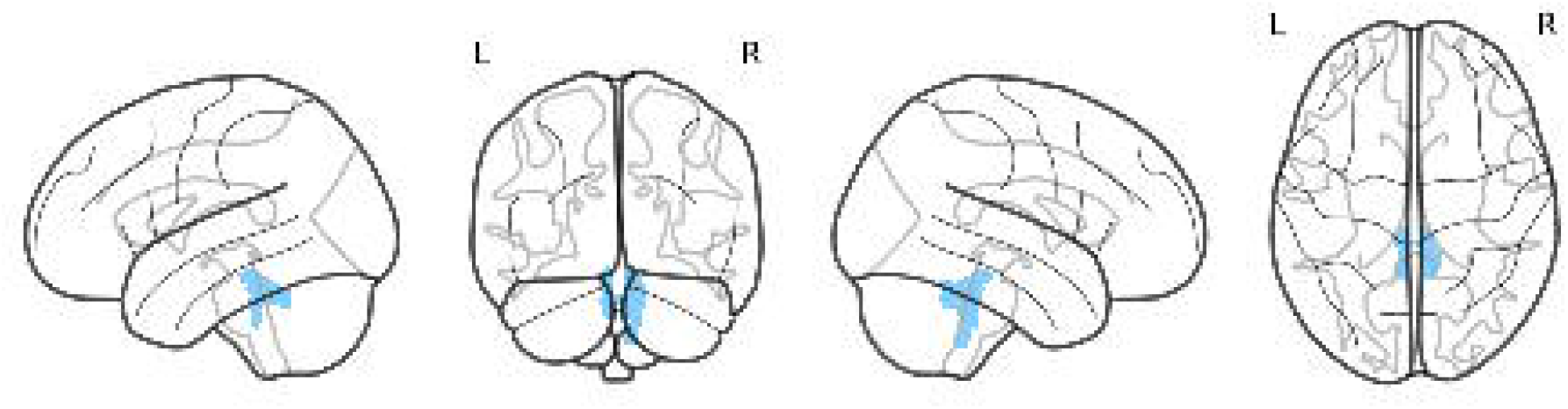

## Discussion

Our main findings are that PSF affects one out of five ischemic stroke patients 3 months post-stroke and that stroke lesions located in motor pathways (more specifically in the corona radiata) and in cerebro-cerebellar tracts may contribute to the occurrence of PSF. Additionally, various predisposing factors may be differently related to facets of the PSF construct, particularly psychological distress. Using a multidimensional tool permitted the identification of clinical, sociodemographic, and neuroradiological determinants related to specific domains of fatigue that have been underexamined in the literature.

In our cohort of non-severe first-ever ischemic stroke patients, 20.8% had MFI-defined PSF. It is substantially lower than the 48% overall and 36% ischemic-subtype prevalence found in the most recent meta-analysis^3^. This is most likely due to the fact that we have excluded patients with severe physical disabilities, with a recent or current mood or anxiety disorder, and our implementation of the MFI instead of the typically used Fatigue Stroke Scale.

Regarding socio-dememographic risk factors, women endorsed greater general fatigue, independent of stroke and psychological distress severity. This finding suggests that specific dimensions of PSF are experienced differently between men and women. Additionally, younger stroke participants endorsed higher mental fatigue. Previous studies have also noted the higher prevalence of neuropsychological impairments, including fatigue, in younger patients^3,18,19^. Researchers suggest that this could reflect how the impact of stroke on daily life may be experienced more abruptly by younger and relatively healthier patients than by those who progressively experience an increase in mental fatigue and a decline in overall health with age. However, since fatigue is measured subjectively, the use of fatigability - an objective index of lower performance due to fatigue - is required to ascertain this hypothesis. Certain facets of PSF were impacted by stroke severity: in particular, patients with higher NIHSS scores reported greater general and physical fatigue. Due to our exclusion of severe stroke patients, the question of whether these are the only dimensions of fatigue influenced by stroke severity remains only partially answered.

Regarding psychological variables, PSF was robustly associated with depression both cross-sectionally and prospectively, in line with the literature^7,16^. Our results also highlight the interplay between anxiety and PSF in the months following stroke - a psychological relationship often neglected in the literature. Nevertheless, since pre-stroke fatigue was not assessed, future studies including it are needed to determine whether anxiety and depression predict PSF independently.

In terms of neurological markers, and in line with the literature, lesion volume was not correlated with PSF^6^. Concerning cerebral integrity and overall PSF, we did not find a significant involvement of the thalamus, or more generally of the brainstem, thus not corroborating the brainstem fatigue generator model^17^. However, our SVR-LSM approach did identify an association between the right corona radiata as well as the right external capsule and overall PSF, in line with some results in the literature^8^. This suggests that lesions in the motor pathways, either in the pyramidal or extrapyramidal system, may be major contributors to PSF^7^. The presence of an association between the severity of fatigue and the lesion location in the corona radiata could also explain the link identified between NIHSS and fatigue.

On the other hand, our network-based approach showed that Mental fatigue and Reduced Activity subdimensions were associated with cerebro-cerebellar networks integrity, which does partially align with the brainstem fatigue generator model. This contrast between voxel and network approaches is not completely unexpected, considering the hypothesis that PSF may be linked to the alteration of white matter tracts underlying brain connectivity and functioning^20,21^. It is essential to acknowledge how examining wide neural connectivity patterns may provide broader perspective than voxel-based analyses.

Our study has notable limitations. First, our sample included minor ischemic stroke patients without severe disability. We believe this reduces how external factors beyond the mechanisms of the stroke interact and influence the state of mental tiredness, as recognized in previous studies^22^. Moreover, since there is evidence that transient ischemic attacks can also result in long-lasting fatigue^23^, restricting our cohort to a more homogeneous stroke population strengthens the validity of our findings. Nevertheless, our criteria impede any generalization of results to patients with hemorrhagic stroke or severe disabilities. Similarly, only cognitive status at inclusion and post-stroke mental fatigue tended to be associated, but the low level of cognitive disability of the study population precludes any firm conclusion on potential associations between post-stroke cognition and PSF. A fine-grained cognitive evaluation and a more diverse sample as in the CADASIL cohort^7^ would be necessary to address this question. Second, the low overlap of lesion location resulting from our moderate sample size limited the SVR-LSM analyses; despite high processing power, the program could not provide convergence beyond 2000 permutations - lower than the default parameters. Third, for the network-based approach,. we chose the Harvard-Oxford atlas over the MNI152 and AAL3 - which are more parcellated - due to the large number of principal components and their incoherence regarding the regions clustered from the latter. Additionally, the Harvard-Oxford atlas does not provide laterality for cortical regions, which may have impacted our findings. Finally, the present study focused on lesion characteristics and psycho-sociodemographic factors, failing to consider the influence of potential biological markers. With this respect, it is important to note that systemic inflammation has been suggested to contribute to PSF, with higher levels of inflammatory biomarkers associated with PSF^3^.

Overall, our study confirms the importance of considering post-stroke fatigue as a multidimensional phenomenon, warranting the use of an appropriate screening tool to capture each of its domains. Clinicians must consider different fatigue profiles among the stroke population depending on factors such as age and sex. Our study also aligns with the fact that mood impairments, such as elevated anxiety and depressivity, are tightly associated with fatigue and further stresses the need to address these difficulties in post-stroke care. White matter alterations were primarily related to levels of fatigue, suggesting that an accumulation of lesions in white matter tracts end up dysregulating brain networks. The ‘effort’ of the brain to compensate for these motor pathways alterations may necessitate an overactivity that is potentially responsible for this syndrome of fatigue. Further examination of brain networks’ disruptions, particularly in white matter, may provide more clarity on this hypothesis.

## Acknowledgments

The authors would like to acknowledge all the participants, Sylvain Ledure for his assistance in conducting the study, and the Clinical Epidemiology Unit of Bordeaux University hospital (USMR), in particular Antoine Bénard and Florence Allais, for their valuable support in the data collection and storage process.

