## Supplementary Material for "Insights from multidimensional analyses of post-stroke fatigue"

### Supplementary Materials

#### *Univariate analyses by groups*

Table 1. Descriptive statistics and comparison between men and women

| | | Men (mean; SD) | Men (%) | Women (mean; SD) | Women (%) | Comparison p,(effect size) | Proportion (χ2; p) |
|---|---|---|---|---|---|---|---|
| Age | | 58.1 (14.1) | / | 55.0 (17.0) | / | 0.176, (0.09) | / |
| Obesity | T1 | / | 12.7 | / | 11.7 | / | 0.040. 1.000 |
| Diabetes | T1 | / | 12.9 | / | 5.9 | / | 2.437; 0.163 |
| Hypertension | T1 | / | 55.2 | / | 38.2 | / | 5.533; **0.021** |
| Dyslipidemia | T1 | / | 83.4 | / | 67.6 | / | 7.157; **0.013** |
| Alcohol consumption | T1 | 0.9 (1.4) | 67.9 | 0.2 (0.4) | 50.0 | **<0.001**, (0.28) | 5.976; **0.017** |
| | T2 | 0.8 (1.1) | / | 0.4 (0.8) | / | 0.310, (0.13) | / |
| HSI | T1 | 1.9 (1.6) | 28.2 | 1.4 (1.4) | 29.4 | 0.433, (0.00) | 0.033; 0.874 |
| | T2 | 0.1 (0.4) | / | 0.0 (0.3) | / | 0.632, (0.04) | / |
| mRS | T2 | 0.7 (0.8) | / | 1.0 (0.8) | / | **0.003**, (0.2) | / |
| NIHSS | T1 | 0.4 (0.8) | / | 0.5 (0.9) | / | 0.938, (0.01) | / |
| | T2 | 0.2 (0.5) | / | 0.2 (0.5) | / | 0.684, (0.03) | / |
| MoCA | T1 | 26.1 (2.7) | / | 26.3 (2.8) | / | 0.496, (0.04) | / |
| | T2 | 26.3 (2.7) | / | 25.1 (5.5) | / | 0.376, (0.06) | / |
| HAD-A | T1 | 5.8 (3.2) | 36.2 | 6.3 (2.7) | 48.5 | 0.127, (0.10) | 3.558; 0.073 |
| | T2 | 4.9 (3.3) | 24.5 | 6.4 (3.6) | 42.6 | **0.002**, (0.20) | 7.944; **0.004** |
| HAD-D | T1 | 2.4 (2.4) | 8.6 | 2.4 (2.2) | 4.4 | 0.861, (0.01) | 1.228; 0.206 |
| | T2 | 3.2 (3.6) | 14.7 | 3.8 (3.2) | 17.6 | **0.047**, (0.13) | 0.365; 0.337 |
| MFI Total | T2 | 43.8 (16.5) | 19.6 | 48.3 (15.5) | 23.5 | **0.036**, (0.10) | 0.457; 0.593 |
| MFI General | T2 | 9.8 (4.3) | 32.5 | 11.8 (4.4) | 50.0 | **0.001**, (0.21) | 6.372; **0.016** |
| MFI Physical | T2 | 9.2 (4.0) | 39.9 | 10.0 (4.3) | 45.6 | 0.207, (0.08) | 0.675; 0.462 |
| MFI Reduced Motivation | T2 | 8.0 (3.5) | 28.8 | 8.3 (3.1) | 36.8 | 0.248, (0.08) | 1.444; 0.274 |
| MFI Reduced Activity | T2 | 9.1 (3.5) | 33.7 | 9.7 (4.2) | 42.3 | 0.354, (0.06) | 1.692; 0.228 |
| MFI Mental | T2 | 7.7 (3.9) | 27.6 | 8.3 (3.6) | 27.9 | 0.112, (0.10) | 0.004; 1.000 |
| Lesion Volume (mm3) | T1 | 3218.5(7489.8) | / | 3206.1(5742.6) | / | 0.971, (0.00) | / |
| Fazekas P | T1 | / | 55.4 | / | 45.5 | / | 1.848; 0.189 |
| Fazekas D | T1 | / | 47.5 | / | 47.1 | / | 0.006; 1.000 |
| Brainstem Leucoencephalopathy | T1 | / | 15.9 | / | 19.1 | / | 0.317; 0.568 |
| Lobar Microbleeds | T1 | / | 9.8 | / | 5.9 | / | 0.826; 0.446 |
| Deep Microbleeds | T1 | / | 10.4 | / | 7.4 | / | 0.435; 0.624 |

*Note: Bold figures indicate significance level at p<0.05*

##### *Univariate analyses – dimensional variables*

Table 2. Spearman correlations between MFI Total and subdimensions scores and the socio-demographic, psychological and neurological variables of interest.

| | MFI Total ρ;p | | MFI General ρ;p | | MFI Physical ρ;p | | MFI Motivation ρ;p | | MFI Activities ρ;p | | MFI Mental ρ;p | |
|---|---|---|---|---|---|---|---|---|---|---|---|---|
| | T1 | T2 | T1 | T2 | T1 | T2 | T1 | T2 | T1 | T2 | T1 | T2 |
| Age | -0.072; 0.274 | | -0.860; 0.194 | | 0.031; 0.640 | | 0.004; 0.954 | | -0.026; 0.696 | | -0.205; **0.002** | |
| MoCA | -0.086; 0.192 | -0.038; 0.565 | 0.033; 0.617 | -0.019; 0.777 | -0.133; 0.044 | -0.063; 0.343 | -0.108; 0.103 | -0.056; 0.402 | -0.126; 0.056 | -0.067; 0.309 | -0.090; 0.176 | 0.012; 0.856 |
| NIHSS | 0.043; 0.513 | 0.135; **0.040** | 0.071; 0.284 | 0.191; **0.004** | 0.090; 0.175 | 0.141; **0.033** | -0.021; 0.748 | 0.048; 0.466 | -0.008; 0.906 | 0.110; 0.086 | 0.063; 0.341 | 0.094; 0.155 |
| HAD-A | 0.219; **<0.001** | 0.480; **<0.001** | 0.164; **0.014** | 0.463; **<0.001** | 0.199; **0.003** | 0.364; **<0.001** | 0.149; **0.026** | 0.344; **<0.001** | 0.165; **0.013** | 0.336; **<0.001** | 0.212; **0.001** | 0.508; **<0.001** |
| HAD-D | 0.403; **<0.001** | 0.720; **<0.001** | 0.327; **<0.001** | 0.642; **<0.001** | 0.390; **<0.001** | 0.590; **<0.001** | 0.338; **<0.001** | 0.638; **<0.001** | 0.352; **<0.001** | 0.587; **<0.001** | 0.283; **<0.001** | 0.546; **<0.001** |
| Lesion Vol. | 0.028; 0.668 | | 0.028; 0.676 | | 0.050; 0.452 | | -0.015; 0.819 | | 0.046; 0.488 | | -0.007; 0.921 | |

*Note: Bold figures indicate significance level at p<0.05*

Table 3. Spearman's correlations between Clinical factors

| | | NIHSS (*ρ*;p) | | MoCA (*ρ*;p) | | HAD Anx. (*ρ*;p) | | HAD Dep. (*ρ*;p) | |
|---|---|---|---|---|---|---|---|---|---|
| | | T1 | T2 | T1 | T2 | T1 | T2 | T1 | T2 |
| Age | | 0.057; 0.380 | -0.026; 0.692 | -0.316; **<0.001** | -0.288; **<0.001** | -0.040; 0.544 | -0.172; **0.009** | 0.125; 0.052 | 0.019; 0.773 |
| NIHSS | T1 | | 0.495; **<0.001** | -0.038; 0.559 | 0.000; 0.997 | -0.044; 0.502 | -0.045; 0.497 | 0.054; 0.407 | 0.035; 0.592 |
| | T2 | | | 0.068; 0.299 | -0.032; 0.625 | -0.027; 0.686 | -0.015; 0.825 | 0.047; 0.472 | 0.036; 0.590 |
| MoCA | T1 | | | | 0.411; **<0.001** | 0.063; 0.338 | -0.066; 0.317 | -0.068; 0.290 | -0.146; **0.027** |
| | T2 | | | | | -0.010; 0.885 | -0.044; 0.506 | -0.006; 0.924 | -0.128; 0.052 |
| HAD-A | T1 | | | | | | 0.485; **<0.001** | 0.255; **<0.001** | 0.170; **0.011** |
| | T2 | | | | | | | 0.171; **0.009** | 0.540; **<0.001** |
| HAD-D | T1 | | | | | | | | 0.423; **<0.001** |

*Note: Bold figures indicate significance level at p<0.05*

***Confounding factors*** were selected according to the table above: every clinical variable that was significantly correlated with another clinical variable associated with an MFI score was included in the following multivariate models as a cofactor.

##### *Multivariate analyses*

Table 4. Multivariate comparison of MFI scores by men and women adjusted for potential confounding clinical factors

| MFI Total ([F]; p) | MFI General ([F]; p) | MFI Physical ([F]; p) | MFI Motivation ([F]; p) | MFI Activity ([F]; p) | MFI Mental ([F]; p) |
|---|---|---|---|---|---|
| [0.369]; 0.544[a] | [4.014]; **0.046**[a] | [0.022]; 0.881[a] | [0.010]; 0.920[a] | [0.014]; 0.907[a] | [0.001]; 0.979[a] |
| [0.708]; 0.401[b] | [6.389]; **0.012**[b] | [0.012]; 0.911[b] | [0.021]; 0.884[b] | [0.087]; 0.769[b] | [0.361]; 0.548[b] |

*Note: Bold figures indicate significance level at p<0.05*
*a - adjusted for HAD Anxiety scores at T2*
*b - adjusted for HAD Depression scores at T2*

Table 5. Correlations between MFI scores and Clinical factors, adjusted for potential confounding Clinical variables

| | | MFI Total (ρ;p) | MFI General (ρ;p) | MFI Physical (ρ;p) | MFI Motiv (ρ;p) | MFI Act (ρ;p) | MFI Mental (ρ;p) |
|---|---|---|---|---|---|---|---|
| Age | | -0.113; 0.089[a] | -0.081; 0.223[a] | -0.022; 0.736[a] | -0.032; 0.632[a] | -0.070; 0.291[a] | -0.267; **<0.001**[a] |
| | | -0.088; 0.185[b] | -0.089; 0.181[b] | 0.001; 0.948[b] | -0.004; 0.948[b] | -0.045; 0.500[b] | -0.226; **<0.001**[b] |
| | | -0.006; 0.926[d] | -0.024; 0.720[d] | 0.066; 0.317[d] | 0.075; 0.258[d] | 0.027; 0.683[d] | -0.175; **0.008**[d] |
| HAD-A | T1 | 0.107; 0.112[e] | 0.090; 0.184[e] | 0.060; 0.376[e] | 0.073; 0.280[e] | 0.085; 0.206[e] | 0.126; 0.061[e] |
| | | 0.155; **0.021**[f] | 0.110; 0.103[f] | 0.087; 0.198[f] | 0.097; 0.149[f] | 0.117; 0.082[f] | 0.147; **0.029**[f] |
| | T2 | 0.520; **<0.001**[g] | 0.471; **<0.001**[g] | 0.387; **<0.001**[g] | 0.425; **<0.001**[g] | 0.387; **<0.001**[g] | 0.514; **<0.001**[g] |
| | | 0.499; **<0.001**[e] | 0.450; **<0.001**[e] | 0.347; **<0.001**[e] | 0.386; **<0.001**[e] | 0.350; **<0.001**[e] | 0.506; **<0.001**[e] |
| | | 0.136; **0.040**[f] | 0.156; **0.018**[f] | 0.041; 0.540[f] | 0.013; 0.849[f] | -0.009; 0.890[f] | 0.269; **<0.001**[f] |
| HAD-D | T1 | 0.385; **<0.001**[c] | 0.302; **<0.001**[c] | 0.342; **<0.001**[c] | 0.339; **<0.001**[c] | 0.351; **<0.001**[c] | 0.270; **<0.001**[c] |
| | | 0.326; **<0.001**[d] | 0.240, **<0.001**[d] | 0.272; **<0.001**[d] | 0.285; **<0.001**[d] | 0.310; **<0.001**[d] | 0.193; **0.003**[d] |
| | T2 | 0.725; **<0.001**[c] | 0.606; **<0.001**[c] | 0.574; **<0.001**[c] | 0.650; **<0.001**[c] | 0.619; **<0.001**[c] | 0.570; **<0.001**[c] |
| | | 0.602; **<0.001**[d] | 0.460; **<0.001**[d] | 0.464; **<0.001**[d] | 0.567; **<0.001**[d] | 0.543; **<0.001**[d] | 0.369; **<0.001**[d] |

*Note: Bold figures indicate significance level at p<0.05*
*a - adjusted for MoCA scores at T1*
*b - adjusted for MoCA scores at T2*
*c - adjusted for HAD Anxiety scores at T1*
*d - adjusted for HAD Anxiety scores at T2*
*e - adjusted for HAD Depression scores at T1*
*f - adjusted for HAD Depression scores at T2*
*g – adjusted for age*

#### Associating lesion location with clinical evaluations

Table 6. List of brain regions belonging to each component produced by the PCA.

| | |
|---|---|
| **GM1** | Temporal Pole; Superior Temporal Gyrus, anterior division; Middle Temporal Gyrus, anterior division; Middle Temporal Gyrus, posterior division; Middle Temporal Gyrus, temporo-occipital part; Inferior Temporal Gyrus, anterior division; Inferior Temporal Gyrus, posterior division; Juxtapositional Lobule Cortex (formerly Supplementary Motor Cortex); Cingulate Gyrus, anterior division; Cingulate Gyrus, posterior division; Herschl’s Gyrus |
| **GM2** | Inferior Temporal Gyrus, temporo-occipital part; Parahippocampal Gyrus, anterior division; Parahippocampal Gyrus, posterior division; Lingual Gyrus; Temporal Fusiform Cortex, anterior division; Temporal Fusiform Cortex, posterior division; Occipital Fusiform Gyrus; Hippocampus (L) |
| **GM3** | Subcallosal Cortex; Caudate (R); Putamen (R); Pallidum (R); Amygdala (R); Accumbens (R) |
| **GM4** | Insular Cortex; Frontal Orbital Cortex; Frontal Opercular Cortex; Central Opercular Cortex; Heschl’s Gyrus (includes H1 and H2); Amygdala (L) |
| **GM5** | Intracalcarine Cortex ; Precuneus Cortex ; Cuneal Cortex ; Lingual Gyrus ; Supracalcarine Cortex ; Occipital Pole |
| **GM6** | Postcentral Gyrus; Superior Parietal Lobule; Supramarginal Gyrus, anterior division; Angular Gyrus; Lateral Occipital Cortex, superior division |
| **GM7** | Middle Temporal Gyrus, temporo-occipital part; Supramarginal Gyrus, posterior division; Angular Gyrus; Parietal Opercular Cortex; Heschl’s Gyrus |
| **GM8** | Caudate (L); Putamen (L); Pallidum (L) |
| **GM9** | Precentral Gyrus; Lateral Occipital Cortex, superior division; Occipital Pole |
| **GM10** | Inferior Frontal Gyrus, pars opercularis; Frontal Opercular Cortex; Amygdala (L); Middle Frontal Gyrus |
| **GM11** | Frontal Pole; Superior Frontal Gyrus; Middle Frontal Gyrus |
| **WM1** | Genu of corpus callosum; Body of corpus callosum; Anterior limb of internal capsule (L); Posterior limb of internal capsule (L); Superior fronto-occipital fasciculus (L) |
| **WM2** | Posterior limb of internal capsule (L); Anterior corona radiata (L); Superior corona radiata (L); External capsule (L); Superior longitudinal fasciculus (L) |
| **WM3** | Splenium of corpus callosum, Posterior thalamic radiation (L); Cingulum (cingulate gyrus) (L) |
| **WM4** | Body of corpus callosum, Anterior limb of internal capsule (R); Posterior limb of internal capsule (R); Superior corona radiata (R); External capsule (R); Superior fronto-occipital fasciculus (R) |
| **WM5** | Medial lemniscus (R); Medial lemniscus (L); Inferior cerebellar peduncle (R), Superior cerebellar peduncle (R); Superior cerebellar peduncle (L) |
| **WM6** | Posterior thalamic radiation (R); External capsule (R); Superior longitudinal fasciculus (R) |
| **WM7** | Posterior thalamic radiation (R); Sagittal stratum (R), Cingulum (cingulate gyrus) (R); Cingulum (hippocampus) (R) |
| **WM8** | Pontine crossing tract; Corticospinal tract (R); Corticospinal tract (L); Medial lemniscus (L), Cerebellar peduncle (R) |
| **WM9** | Retrolenticular part of internal capsule (R); Posterior corona radiata (R), Ucinate fasciculus (R) |
| **WM10** | Retrolenticular part of internal capsule (L); Posterior corona radiata (L) |
| **WM11** | Middle cerebellar peduncle; Corticospinal tract (R); Inferior cerebellar peduncle (L); Superior cerebellar peduncle (L) |
| **WM12** | Sagittal striatum (L); Cingulum (hippocampus) (L) |

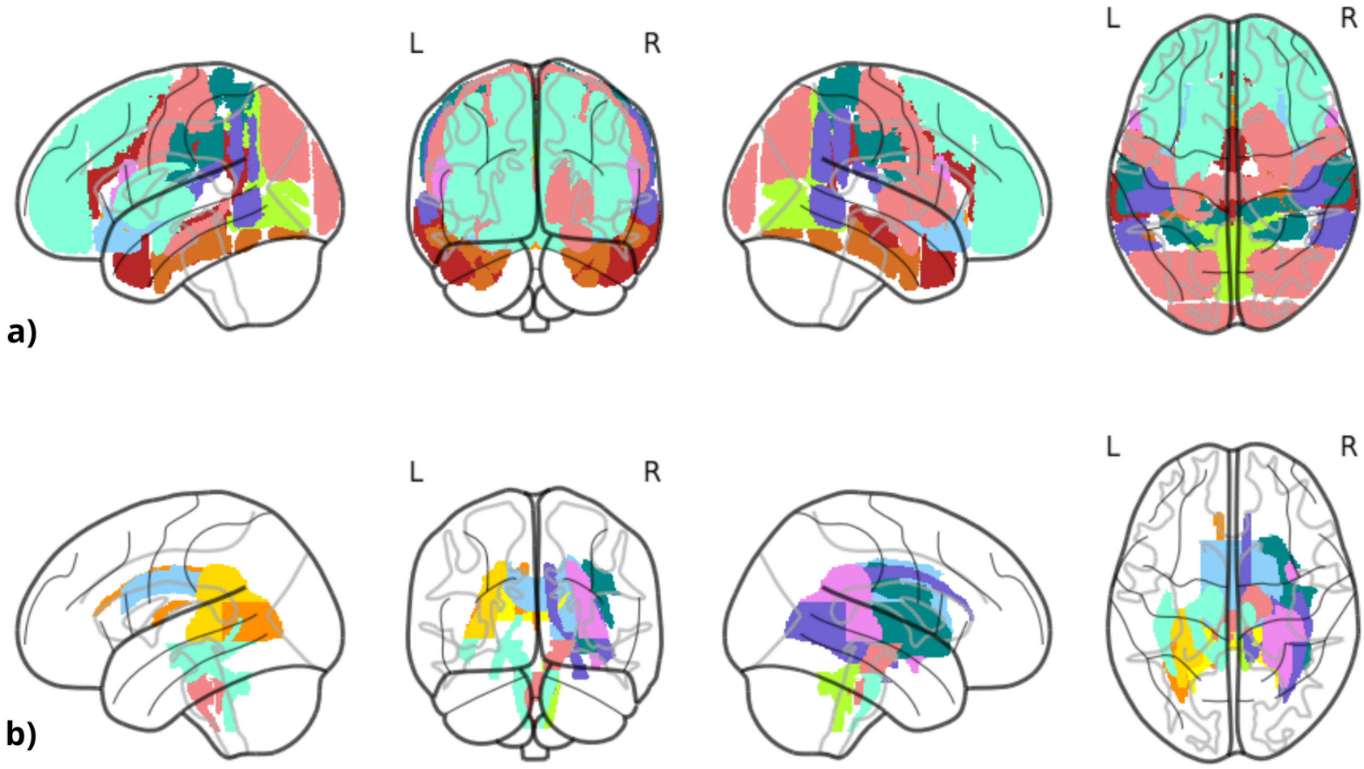

Figure 1. All components obtained from the PCA approach.

*a) Gray matter (GM) components: GM1 in gold, GM2 in red, GM3 in chocolate, GM4 in dark orange, GM5 in light blue, GM6 in green, GM7 in teal, GM8 in purple, GM9 in coral, GM10 in pink, GM11 in turquoise.*

*b) White matter (WM) components: WM1 in yellow, WM2 in red, WM3 in chocolate, WM4 in dark orange, WM5 in light blue, WM6 in green, WM7 in teal, WM8 in purple, WM9 in coral, WM10 in pink, WM11 in gold, WM12 in turquoise.*
